# Examining the Role of Extrachromosomal DNA in Lung Cancer

**DOI:** 10.1101/2025.06.03.657117

**Authors:** Azhar Khandekar, Phuc H. Hoang, Jens Luebeck, Marcos Díaz-Gay, Wei Zhao, John P. McElderry, Caleb Hartman, Mona Miraftab, Olivia W. Lee, Kara Barnao, Kristine Jones, Amy Hutchinson, Belynda Hicks, Erik Bergstrom, Yang Yang, Martin A. Nowak, Nathaniel Rothman, Angela C Pesatori, Dario Consonni, Robert Homer, Marina K. Baine, Lynette M. Sholl, Philippe Joubert, Charles Leduc, William D. Travis, Soo-Ryum Yang, Qing Lan, David C. Wedge, Lixing Yang, Stephen J. Chanock, Tongwu Zhang, Ludmil B. Alexandrov, Maria Teresa Landi

## Abstract

The role of extrachromosomal DNA (ecDNA) in lung cancer, particularly in subjects who never smoked (LCINS), remains unclear. Examination of over 1200 whole-genome-sequenced lung cancers identified ecDNA in 18.9% of patients. Recurrent amplification of *MDM2* and other oncogenes *via* ecDNA possibly drives a LCINS subset. Tumors harboring ecDNA showed worse overall survival than tumors harboring other focal amplifications. A strong association with whole-genome doubling suggests most ecDNA reflects genomic instability in treatment-naïve lung cancer.

## MAIN

Extrachromosomal DNA (ecDNA) is a potent mechanism for oncogene amplification in human cancer^1^, exhibiting unique characteristics such as a circular structure^2^, non-Mendelian inheritance^3,4^, and an altered epigenetic and transcriptional landscape^5^. Prior studies have linked ecDNA to worse overall survival in pan-cancer analyses, highlighting its clinical significance^6–8^ and potential as a therapeutic target^9^. Recent advances in computational techniques^10^ have enabled large-scale detection of ecDNA from whole-genome sequencing (WGS) data. However, only three pan-cancer studies have examined ecDNA in primary non-small cell lung cancer cohorts with over 100 samples, focusing predominantly on patients of European descent^6–8^. Two studies^6,7^ analyzed lung cancer samples from the United States, reporting ecDNA prevalence rates of about 21% in 36 lung adenocarcinomas (LUAD) and 27% in 47 lung squamous cell carcinomas (LUSC). Additionally, a UK lung cancer study^8^ analyzed 14,778 patients across 39 tumor types, including 718 LUAD tumors and 378 LUSC tumors, detecting ecDNA in 8.2% of LUAD and 22.4% of LUSC cases. All three studies^6–8^ reported lung cancer findings as part of broader pan-cancer analyses and did not distinguish between lung cancers in subjects who have never-smoked (LCINS) and subjects who have smoked (LCSS). Notably, based on the high prevalence of the tobacco-associated^11^ mutational signature SBS4 in these studies, more than 84% of the analyzed lung cancer cases were from individuals with a history of smoking.

Despite these prior studies, the clinical relevance of ecDNA in lung cancer remains unclear, including its prevalence across histologies, geographical areas, genetic ancestries, and biological sexes. Its role in LCINS, LCSS, and subjects who were exposed to passive smoking also requires further exploration. To address these questions, here, we analyzed WGS data from 1,216 treatment-naïve lung cancers from diverse patient populations (**Fig. 1*a***; **Supplementary Fig. 1**). This large international cohort enabled a comprehensive evaluation of ecDNA’s association with genomic alterations, as well as its prevalence across smoking status, biological sexes, ancestries, and 18 areas defined primarily by country of residence. The dataset includes 871 LCINS, sourced from the Sherlock-Lung^12^ (*n*=817), the Environment And Genetics in Lung Cancer Etiology (EAGLE^13^, *n*=25), the Cancer Genome Atlas (TCGA^14^, *n*=15), and other publicly available cohorts (*n*=14)^15–19^, representing patients from 17 geographical areas (**Supplementary Fig. 2*a***). Additionally, it comprises 345 LCSS, drawn from EAGLE^13^ (*n*=221) and TCGA^14^ (*n*=83) and other publicly available cohorts (*n*=41)^15–19^, spanning five geographical areas (**Supplementary Fig. 2*b***). The majority of tumors were LUAD (*n*=1,023; 84.1%) or LUSC (*n*=67; 5.5%), with carcinoids (*n*=61; 5%, exclusively in LCINS) and other rarer histological lung cancer subtypes (*n*=65; 5.3%) also represented. Amongst Sherlock-*Lung* LCINS, 250 were exposed to secondhand tobacco smoke, while 208 had documented non-exposure (**Fig. 1*a***).

**Figure 1:**
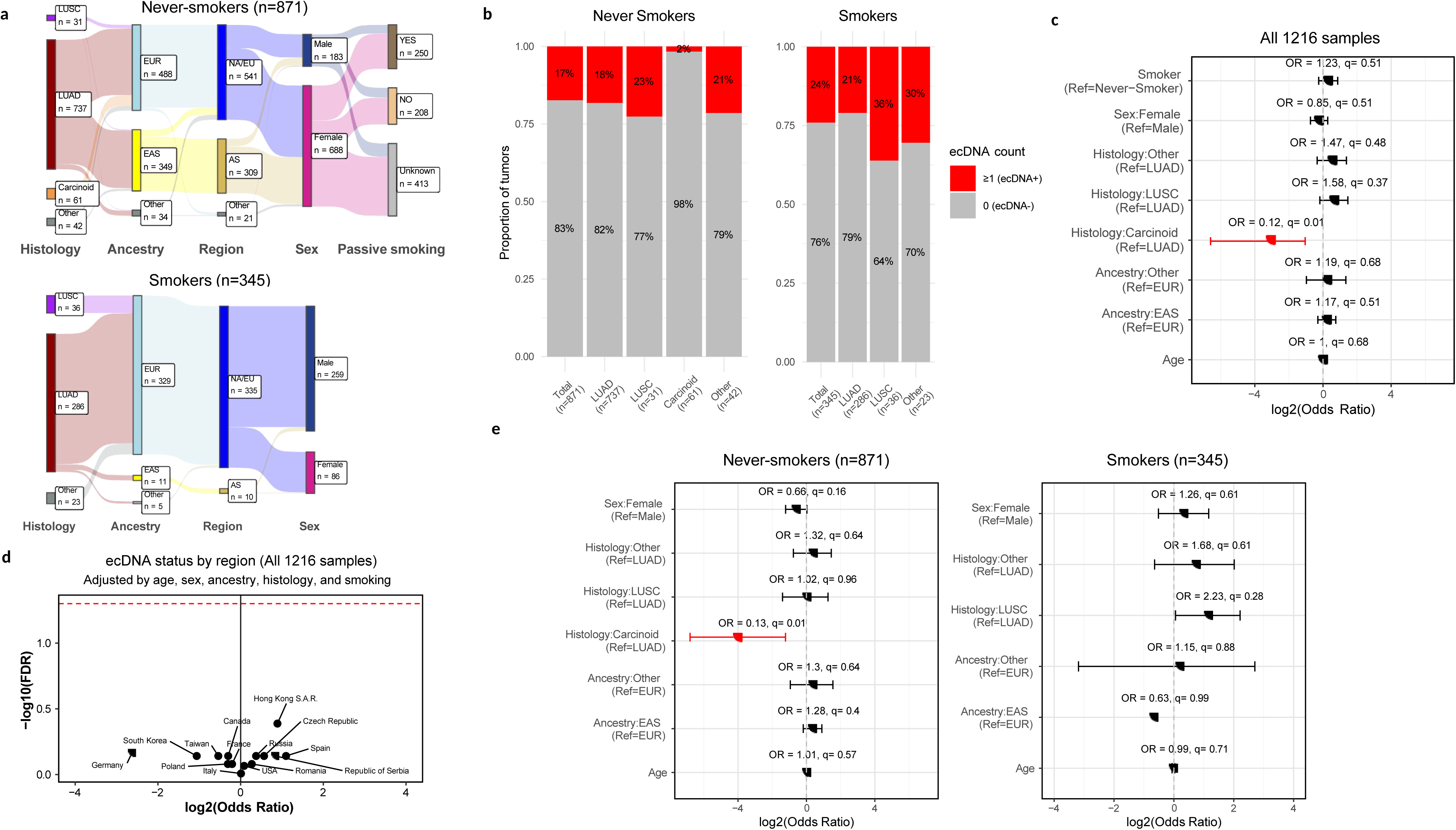
Landscape of extrachromosomal DNA in lung cancer across histologies, sex, ancestry, countries, and smoking status. ***a)*** Epidemiological features of the LCINS (top) and LCSS (bottom) cohorts. Total number of samples in each category is indicated. ***b)*** The number and proportion of tumors harboring ecDNA among LCINS and LCSS, stratified by histology. Samples with 1 or more ecDNA were classified as ecDNA^+^, and samples with no ecDNA were classified as ecDNA^-^. ***c)*** Forest plot of the full cohort (LCINS and LCSS) from a multivariate logistic regression analysis of key epidemiological and clinical features and the presence of ecDNA in the tumors. ***d)*** Volcano plot showing log_2_ odds ratio (x-axis) and log_10_ FDR corrected q-value (y-axis) for a one-vs-all statistical comparison of ecDNA prevalence by region. ***e)*** Forest plots for multivariate logistic regression models testing key epidemiological and clinical features with the presence of ecDNA in the tumors, stratified by smoking status. NA/EU: North America or Europe region, AS: Asia region EAS: East Asian genetic ancestry super-sample, EUR: European genetic ancestry super-sample, LUAD: lung adenocarcinoma, LUSC: lung squamous cell carcinoma.

To detect and reconstruct ecDNA across the cohort, a computational pipeline that accounts for both tumor purity and ploidy was developed (**Methods**). A total of 231 samples harbored at least one ecDNA (18.9%), with similar prevalence between LCINS (17%) and LCSS (23%; q-value: 0.21; **Fig. 1*b***). No statistically significant differences were observed in the prevalence of ecDNA across various histological subtypes, biological sexes, ancestries or age at diagnosis (**Fig. 1*c*&e**), except for carcinoids, which showed a lower prevalence compared to LUAD in the overall analysis (OR=0.12; q=0.01) and when LCINS were considered separately (OR=0.13; q=0.01). ecDNA was present at similar rates across all geographic locations (**Fig. 1*d***; **Supplementary Fig. 2*c-d***), as indicated by a one-versus-all multivariate logistic regression for each country.

There was also no difference in ecDNA prevalence between LCSS and LCINS (**Fig.1*c***). ecDNA was not associated with tumor stage (**Supplementary Fig. 3*a&b***), subsequent development of metastases (**Supplementary Fig. 3*c&d***) or passive smoking status (**Supplementary Fig. 3*e***). These findings remained consistent when analyses were restricted to only LUAD cases in either LCSS or LCINS (**Supplementary Fig. 4**).

To examine the association between ecDNA and genomic features previously linked to genome instability^20–23^, we evaluated the relationship of ecDNA with whole-genome doubling (WGD), chromothripsis, tumor mutational burden, mitochondrial copy number, overall genome instability index (wGII), and telomere length tumor/normal ratio. Amongst these, only WGD showed a significant association with ecDNA in both LCSS and LCINS (**Fig. 2*a***), as determined by logistic regression adjusted for age, sex, ancestry, histology, and tumor purity. Specifically, ecDNA was present in 7.1% of LCINS and 8.9% of LCSS without WGD, but its prevalence increased markedly in WGD-positive tumors from LCINS (24.9%, OR=4.0; q=1.24 x 10^-8^) and LCSS (30.3%, OR=5.85; q=8.06 x 10^-4^). The same association was observed when only LUAD was examined (**Supplementary Fig. 5*a***). Moreover, this association was further validated in a pan-cancer dataset of 974 samples with high-confidence WGD annotation^14^, using logistic regression adjusted for age, purity, sex, and cancer type (OR=2.81; q=1.63 x 10^-6^; **Supplementary Fig. 5*b***).

**Figure 2.**
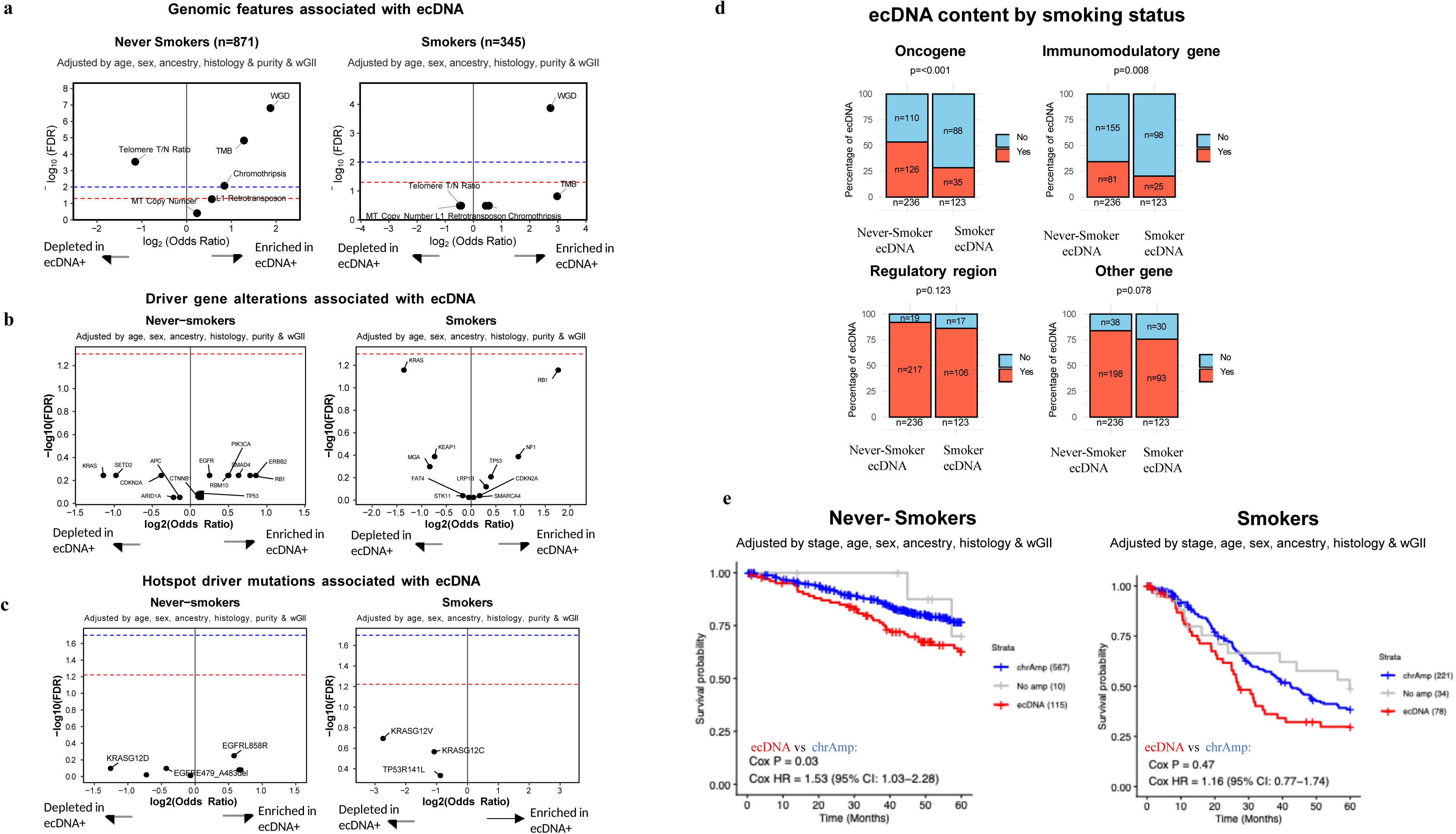
Genomic and clinical characteristics associated with ecDNA. ***a)*** Volcano plot of logistic regression models of genomic features in association with ecDNA status for LCINS (left) and LCSS (right). ***b)*** Volcano plot of a logistic regression model of driver gene alterations in a sample in association with ecDNA status for LCINS (left) and LCSS (right). ***c)*** Volcano plot of a logistic regression model of hotspot driver mutations in a sample in association with ecDNA status for LCINS (left) and LCSS (right). In all volcano plots, the x-axes reflect the log_2_ odds ratio, and the y-axes correspond to the log_10_ FDR q-value. An FDR q-value threshold of 0.05 is indicated with the dashed red line, an FDR q-value threshold of 0.01 is indicated with the dashed blue line. ***d)*** Bar plot indicating the proportion of ecDNA containing a particular genomic element in LCINS vs LCSS. ***e)*** Kaplan–Meier survival curves for 5-year overall survival stratified by the mode of amplification status for lung cancers in LCINS (left) and LCSS (right). P-values for significance and HRs of the difference were calculated using two-sided Cox proportional-hazards regression with adjustment for age, sex, tumor stage, ancestry and histology and are indicated within each plot.

In addition to WGD, chromothripsis (OR=1.76; q=0.048; **Fig. 2*a***) and telomere length tumor/normal ratio, (OR=1.76; q=0.01; **Fig. 2*a***) were also significantly associated with ecDNA, but this was exclusively observed in LCINS. Prior studies have indicated that chromothripsis may contribute to the formation of certain ecDNA^23^. In this study, 115 out of 151 LCINS with ecDNA (76.2%) and 60 out of 80 LCSS with ecDNA (75%) exhibited chromothripsis (**Supplementary Tables 1-2**). Nevertheless, in samples where both ecDNA and chromothripsis were detected, overlap between ecDNA segments and chromothripsis regions was rare, occurring more frequently in LCINS—observed in 31 of 115 LCINS cases compared to 4 of 60 LCSS (OR=5.01, p=0.02 comparing LCINS *vs.* LCSS; **Supplementary Fig. 5*c***). This suggests that there could be potentially different mechanisms of ecDNA formation between LCINS and LCSS. However, while the majority of ecDNA emerges in genomically unstable lung cancers that also exhibit chromothripsis, the actual genomic overlap between ecDNA and chromothripsis regions is limited in both lung cancer types. The association between shortened telomere length and ecDNA persisted even when using an alternative telomere estimation method^24^ (**Supplementary Fig. 5d)**, and whether it was input as a binary or numeric variable (**Supplementary Fig. 5e**). This result is consistent with prior studies that have reported that telomere length is linked with cell division^25^ and that cells with higher ecDNA levels divide faster^26^.

To evaluate the relationship between ecDNA presence and lung cancer driver genes, we analyzed driver genes altered in at least 10 samples as well as driver hotspot mutations occurring at the same genomic position in at least 10 samples. Using multivariate logistic regression, adjusted for age, sex, tumor purity, ancestry, and histology, no significant associations were found between ecDNA and individual driver gene alterations (**Fig. 2*b***) or between ecDNA and individual driver alterations (**Fig. 2*c***).

Mutational signature analysis provides insights into both the endogenous and exogenous mutational processes that have been active throughout the lineage of a cancer genome^11^. We investigated the association between ecDNA presence and mutational signatures. As expected^27^, multiple copy-number and structural variant signatures were associated with ecDNA presence similarly in LCSS and LCINS (**Supplementary Fig. 6*a-b***). Additionally, in LCINS, ecDNA-positive samples were enriched for clock-like signatures (SBS1, SBS5, ID1, ID2) and, as previously reported^19,28^, APOBEC-associated signatures (SBS2, SBS13; **Supplementary Fig. 6*a***). In LCSS, enrichment was observed only for SBS3 (**Supplementary Fig. 6*b***), a signature previously associated with homologous recombination deficiency^29^.

Previous studies have shown that ecDNA can harbor oncogenes, regulatory elements, and immunomodulatory genes in varying configurations^3,8,30,31^. In our cohort, 151 (17%) LCINS and 80 (23%) LCSS harbored ecDNA, with a total of 236 ecDNAs in LCINS and 123 in LCSS, reflecting multiple ecDNAs in some tumors. Among these, ecDNA in LCINS were enriched with oncogenes (p<0.001) and immunomodulatory genes (p=0.008) compared to LCSS (**Fig. 2*d***; **Supplementary Fig. 7*a****-**b***). Further, ecDNA containing oncogenes exhibited elevated copy numbers (CN) compared to other ecDNA without oncogenes in LCINS (p=1.5 x 10^-3^) and LCSS (5.8 x 10^-3^), suggesting greater selective pressure for these ecDNAs (**Supplementary Fig. 7*c***), as previously reported in pan-cancer^8^. *MDM2* was the most frequently amplified oncogene on ecDNA in both LCINS (*n*=36) and LCSS (*n*=5). Given its prominence, we examined the relationship between ecDNA-driven *MDM2* amplification and *TP53* alterations, as MDM2 negatively regulates TP53 through ubiquitination^32^. We found that *MDM2* amplification on ecDNA was mutually exclusive with *TP53* alterations (OR=7.51, p=0.01). In LCINS, the next four most prevalent oncogenes that did not co-occur with *MDM2* were *TERT* (n=10), *CCND1* (*n*=8), *MYC* (*n*=8), and *EGFR* (*n*=7; **Supplementary Table 3**). In LCSS, no oncogene besides *MDM2* was amplified on ecDNA in more than two samples (**Supplementary Table 4).**

Prior pan-cancer studies have shown that cancers harboring ecDNA amplifications exhibit worse overall survival^6–8^. To evaluate this association in the lung cancer cohort, survival analysis was conducted on 692 LCINS and 333 LCSS with available stage and covariate data. Lung cancers were categorized as harboring ecDNA, other focal amplifications, or no focal amplifications. Multivariate Cox proportional-hazards regression analysis revealed that patients with ecDNA-positive lung cancers had worse overall survival than those with only chromosomal amplifications; this association was statistically significant in LCINS (hazard ratio=1.53; p=0.03; **Fig. 2*e***).

In summary, we examined the full spectrum of ecDNA in the largest dataset of whole-genome sequenced lung cancers to date, from many geographical regions and ancestry, and with unique information on tobacco smoking status. ecDNA was present in 17% of LCINS and 23% of LCSS. ecDNA in LCINS were enriched with oncogenes and immunomodulatory genes in comparison to LCSS. In LCINS, the *MDM2* locus was often amplified through ecDNA, and shortened telomere length was associated with ecDNA. No associations were identified between presence of ecDNA and biological sexes, smoking status, or ancestries. The carcinoid subtype was depleted of ecDNA; no other differences by histology were observed. However, tumors carrying ecDNA showed worse overall survival in comparison to those with focal chromosomal amplification, particularly in LCINS. A strong association was observed between ecDNA and WGD, and this observation was validated in an independent pan-cancer cohort, suggesting that most ecDNA in treatment-naïve lung cancer are likely to be a product of genomic instability. Notably, ecDNA carrying oncogenes could contribute to driving LCINS and affecting prognosis. Future studies will be needed to dissect the mechanisms of ecDNA formation and maintenance in lung cancer and evaluate its clinical implications in both treatment-naïve and post-treatment settings.

## METHODS

### Criteria for sample inclusion

Similar to our previous publication^33,34^, we used seven rigorous criteria for sample inclusion, as follows. (1) Sequencing coverage. We maintained a minimum average sequencing coverage of more than 40× for tumor samples and more than 25× for normal samples. (2) Contamination and relatedness. Cross-sample contamination was limited to less than 1% by Conpair^35^, and detected relatedness was maintained at less than 0.2 by Somalier^36^. (3) Copy-number analysis. Individuals with abnormal copy-number profiles in normal samples were excluded, as determined by Battenberg^37^. (4) Mutational signatures. Tumor samples exhibiting mutational signatures SBS7 (associated with exposure to ultraviolet light^11^) and SBS31 (associated with platinum chemotherapy)^38^ were removed from the analysis. (5) Tumor type validation. Tumor samples reported as non-lung cancer or not originating from primary lung cancer were excluded. (6) WGS quality control. Tumor samples with a total genomic alteration count of less than 100, or samples with a total genomic alteration less than 1,000 and a number of reads per clonal copy (NRPCC) less than 10, were excluded as low-quality samples. (7) Multiple-region sequencing. In rare cases in which multiple regions of a tumor were sequenced, only one high-purity tumor sample was included to avoid redundancy. These stringent criteria were applied consistently to ensure the robustness and reliability of the data collected for the Sherlock-Lung study. Following the application of these criteria, a total of 1217 samples were available for analysis, but one sample with unknown smoking status was excluded, resulting in a total of 1216 samples across the cohort.

### Determination of sample purity and ploidy

The Battenberg algorithm (v.2.2.9)^37^ was used to determine the sample purity and ploidy. Any somatic copy number variations (CNV) profile determined to have low-quality after manual inspection underwent a refitting process. This process required new tumor purity and ploidy inputs, either estimated by Ccube (v.1.0)^39^ or recalculated from local copy number status. The Battenberg refitting procedures were iteratively executed until the final CNV profile was established and met the criteria of manual validation check.

### ecDNA detection and characterization

CNVKit v.0.9.6^40^ was run in tumor-normal mode to call somatic CNVs against the matched normal whole-genome sequenced samples for each patient, using the log ratio of tumor to normal read depths (logR) and the circular binary segmentation (CBS) method. Note that this method, by default, assumes a sample purity of 1 and a sample ploidy of 2 as it does not correct for purity and ploidy. This limitation can lead to high copy number segments being missed due to low purity^41^. Due to the wide range of tumor purities in our dataset, consistent with prior reports in lung cancer^41^, we rescaled the total copy number estimate using the purity and ploidy estimated from Battenberg according to the following equation:

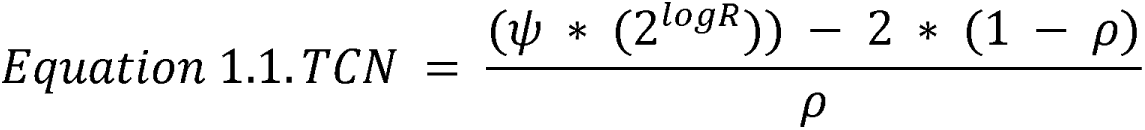

where ψ is the tumor ploidy, ρ is the tumor purity, and logR represents the log ratio of the tumor read depth to the normal read depth (**Supplementary Fig. 8a**). These copy number calls were used to identify putative ‘seed’ regions. We then used AmpliconArchitect (v.1.3)^10^ to reconstruct the architecture of the amplicons and AmpliconClassifier (v.1.3.1) to classify amplicons according to the most probable mechanism of formation (ecDNA, BFB, complex non-cyclic, or linear). Visual inspection and verification were performed for all ecDNA amplicon regions. This pipeline allowed for samples with low purities to be utilized as it completely removed the effect of tumor purity in ecDNA detection (**Supplementary Fig. 8b**). As a result, more ecDNA^+^ samples were detected (**Supplementary Fig. 8b**) and all 1216 samples could be used for ecDNA analysis (**Supplementary Fig. 8c**). Of the additional ecDNA detected, eight of these were *EGFR*-ecDNA (seven in LCINS and one in LCSS) and each of these ecDNA except for one also harbored a driver mutation (**Supplementary Table 5**). Among the eight EGFR-ecDNA+ samples, five had RNA-seq data, which showed very high EGFR expression (**Supplementary Fig. 8d**).

### Determination of whole genome doubling status

As previously done^42^, samples were considered to have a whole genome doubling event if the major copy number was ≥3 for more than 50% of copy number segments.

### Chromothripsis detection and characterization

To detect chromothripsis regions in genomes, we applied the computational algorithm ShatterSeek^43^, which employs a set of statistical criteria to identify chromothripsis given an input of copy number and structural variants. We used the most stringent criteria corresponding to high confidence chromothripsis calls. Specifically, these criteria were: *(i)* a cluster of structural variants (>6 DUP/DEL/h2hINV/t2tINV), *(ii)* oscillating CNV between two states (>7 CNV events), *(iii)* chromosomal enrichment and distribution of DNA breakpoints (p-value<0.05), *(iv)* randomness of fragment joins (p-value>0.05) and/or ≥4 inter-chromosomal rearrangements between multiple chromosomes. The input CNV calls were generated using CNVkit in tumor-normal mode with default parameters, and the input structural variants calls were generated using a union of Meerkat (v.0.189) and Manta (v.1.6.0) calls with the recommended filtering.

### Association of ecDNA with demographic and clinical features

Demographic and clinical features where information was available for all samples (age, sex, ancestry, and tumor histology and purity) served as input variables in a multivariate logistic regression model with ecDNA presence or absence as the dependent outcome variable, and these models were constructed separately for LCINS and LCSS. In addition, separate models for passive smoking, tumor stage, and presence of metastases were run along with the rest of the variables in both LCINS and LCSS.

### Determination of telomere length tumor/normal lung tissue ratio

We estimated telomere length (TL) in kilobases using TelSeq (v.0.0.2)^44^. As was previously done^34^, we used seven as the threshold for the number of TTAGGG/CCCTAA repeats in a read for the read to be considered telomeric. The TelSeq calculation was done individually for each read group within a sample, and the total number of reads in each read group was used as weight to calculate the average TL for each sample. The telomere length ratio (log_2_ scale) between tumor and normal tissue was then calculated for each patient and served as the independent variable in multivariable logistic regressions.

### Association of mutational signatures and ecDNA status

Mutational signature attributions were determined as was previously done, based on the methodology in Díaz-Gay et. al^34^. SigProfilerMatrixGenerator^45,46^ was used to create the mutational matrix for all types of somatic mutations. The deconvolution of mutational signatures was performed by SigProfilerExtractor (v.1.1.3)^47^ using the whole-genome sequencing setting and the COSMIC mutational signatures (v3.2) as reference. Signature extractions were performed separately for LCINS and LCSS. Signature attributions were input into a multivariate logistic regression model as either present or absent if the signature was present in less than 50% of the samples. If the signature was present in ≥50% samples (*e.g.*, SBS1 and SBS5), it was input as either above median or below median. For SBS1 and SBS5, the signature attributions were also input as continuous numeric variables to confirm the significant associations found with inputting them as above median or below median.

### Association of genomic features with ecDNA status

Multivariate logistic regression was performed for each feature individually, with adjustment for covariates (age, sex, ancestry, histology, wGII, and tumor purity). All continuous numeric variables were both input as raw numeric values and binarized and input as above median/below median.

### Association of driver genes and ecDNA status

The IntOGen pipeline (v.2020.02.0123)^48^, which combines seven state-of-the-art computational methods, was employed with default parameters to detect signals of positive selection in the mutational patterns of driver genes across the cohort. The status of each driver gene (altered or unaltered) was then input into a logistic regression model with covariates age, sex, ancestry, histology, wGII, and tumor purity, and ecDNA status was the output. These sets of genes were considered altered if there was a focal deletion overlapping the gene, an inactivating (stop or missense) mutation, or an activating gain of function mutation.

### Association of driver mutations and ecDNA status

To identify driver mutations within the set of identified driver genes, we implemented a rigorous and multifaceted strategy, considering multiple criteria: *(i)* the presence of truncating mutations specifically in genes annotated as tumor suppressors, *(ii)* the recurrence of missense mutations in a minimum of 3 samples, *(iii)* mutations designated as “Likely drivers” with a boostDM score^49^ exceeding 0.5, *(iv)* mutations categorized as either “Oncogenic” or “Likely Oncogenic” based on the criteria established by OncoKB^50^, an expert-guided precision oncology knowledge base, *(v)* mutations previously recognized as drivers in the TCGA MC3 drivers paper^51^, and *(vi)* missense mutations characterized as “likely pathogenic” in genes that are annotated as tumor suppressors, as described in Cheng et al.^52^ Any mutation meeting one or more of these criteria was recognized as a potential driver mutation.

### Calculation of genome instability index score

To calculate the genome instability index score (wGII), the method devised in Bailey et al.^8^ was used. The overall ploidy was calculated as the length-weighted average of all copy number segments. The proportion of segments that were aberrant (where total copy number was greater than the ploidy or less than the ploidy) was computed for each chromosome, these proportions were summed, and the total was divided by 22.

### Identifying regulatory regions on ecDNA

A set of lung enhancer regions were downloaded from EnhancerAtlas^53^, and each ecDNA region was overlapped with these enhancer regions with the requirement that the entire enhancer had to be fully contained (100% overlap) on the ecDNA. A set of lung cancer promoter regions was extracted using a R script which downloaded regions 1000 base pairs upstream of transcription start sites for all genes.

### Defining a set of immunomodulatory genes

ecDNAs were considered immunomodulatory if a gene from the significant gene sets mapped to an ecDNA that did not contain an oncogene, and that significant gene set had an immunomodulatory function (GO terms: 0006968, 0002228, 0042267, 0001906, 0001909, 0002698, 0001910, 0031341, 0002367, 0002695, 0050866, 0051250, 0050777), following the procedure in Bailey et. al^8^. This resulted in a set of 376 genes.

### Mitochondrial copy number estimation

Mitochondrial copy number was estimated per sample using the ratio of mitochondrial read depth to nuclear genome depth, adjusted for tumor ploidy and purity, following the procedure in Reznik *et al.*^54^

### Identification of LINE-1 retrotransposons

To identify putative transposable elements (TE), we utilized a pipeline called TraFiC-mem (v.1.1.0)^55^ (Transposome Finder in Cancer) retaining only TE insertions that passed the default filtering criteria. Detailed methods can be found in Zhang et. al^42^.

## Statistical analysis

All statistical analyses and graphic displays were performed using the R software v4.2.3 1011 (https://www.r-project.org/). Standard logistic regression with adjustment for covariates was used for the enrichment analyses of categorical variables. We applied Firth’s penalized logistic regression using the logistf package in R to model the association between ecDNA status and geographic regions, in order to reduce small-sample bias and address potential issues of data separation. For the comparison of numerical variables across groups, we used non-parametric Mann-Whitney (Wilcoxon rank sum) tests. P-values <0.05 were considered statistically significant. Survival analyses were performed using Cox proportional-hazards regression adjusted for age at diagnosis, sex, tumor stage, tumor histology, ancestry, and wGII. If multiple hypothesis testing was required, we used a false-discovery rate correction based on the Benjamini-Hochberg method and reported FDR. FDR <0.05 were considered statistically significant.

## SUPPLEMENTARY FIGURE LEGENDS

**Supplementary Fig 1. Additional cohort information.** Sankey diagrams showing the numbers of samples with clinical variables (stage, metastasis) as well as survival data available for ***a)*** LCINS and ***b)*** LCSS

**Supplementary Fig 2. Distribution of ecDNA harboring lung cancers in LCINS and LCSS by country.** Maps showing the worldwide prevalence of lung cancers harboring ecDNA (ecDNA^+^) by country for the ***a)*** LCINS cohort and ***b)*** LCSS cohort. Volcano plot of a multivariate one-vs-all logistic regression model for each country, with ecDNA status as the outcome, for ***c)*** LCINS and ***d)*** LCSS.

**Supplementary Fig 3. Analysis of the association between stage, metastasis, and passive smoking with ecDNA status in all LCINS and LCSS.** Forest plots of logistic regression model with ecDNA status as the outcome and the variables for stage in **a)** LCINS and **b)** LCSS; metastasis in **c)** LCINS and **d)** LCSS; and passive smoking in **e)** LCINS. The total number of samples with information for all variables is indicated above each plot.

**Supplementary Fig 4. Analysis of the association between stage, metastasis, and passive smoking with ecDNA status in lung adenocarcinomas**. Forest plots of logistic regression model with ecDNA status as the outcome and the variables for stage in **a)** LCINS and **b)** LCSS; metastasis in **c)** LCINS and **d)** LCSS; and passive smoking in **e)** LCINS. The total number of samples with information for all variables is indicated above each plot.

**Supplementary Fig 5. Genomic features associated with the presence of ecDNA. *a)*** Volcano plot of logistic regression models of genomic features in association with ecDNA status for lung adenocarcinoma in LCINS (left) and lung adenocarcinoma in LCSS (right). In all volcano plots, the x-axes reflect the log_2_ odds ratio, and the y-axes correspond to the log_10_ FDR q-value. An FDR q-value threshold of 0.05 is indicated with the dashed red line, an FDR q-value threshold of 0.01 is indicated with the dashed blue line. ***b)*** Forest plot of multivariate logistic regression modeling the presence or absence of ecDNA in relation to whole-genome doubling in a pancancer dataset comprising four different tumor types from the PCAWG project, adjusted for the covariates of age, purity, sex, and tissue type. All significant associations are indicated in red. ***c)*** Bar plot indicating the proportion of samples that contained overlapping chromothripsis and ecDNA regions (red) and the proportion of all samples that did not contain any overlap (gray) for all samples that had both an ecDNA and chromothripsis event detected. ***d)*** Forest plot of multivariate logistic regression modeling the presence or absence of ecDNA in relation to telomere length estimates from Telhunter, whereby the telomere length ratio (tumor/normal) was input as a continuous numeric variable. **e)** Boxplot indicating the distributions and median telomere length ratio (tumor/normal) for ecDNA+ vs ecDNA-samples, both in never-smokers (left) and smokers (right). The p-value from a multivariate logistic regression with age, gender, tumor purity, histology, ancestry and wGII score is also indicated.

**Supplementary Fig 6. Association of mutational signatures with ecDNA status.** Volcano plots of logistic regression models of the activities of Single base substitution (SBS), Indel (ID), copy number (CN) and structural variant (SV) signatures in association with ecDNA in lung cancers of ***a)*** LCINS and ***b)*** LCSS. The logistic regression models were adjusted for age, sex, ancestry, histology, and purity. In all volcano plots, the x-axes reflect the log_2_ odds ratio, and the y-axes correspond to the log_10_ FDR q-value. An FDR q-value threshold of 0.05 is indicated with the dashed red line, an FDR q-value threshold of 0.01 is indicated with the dashed blue line.

**Supplementary Fig 7. Genomic content of ecDNA**

**a)** Upset plot indicating the number of ecDNA with oncogenes, immunomodulatory genes, promoter or enhancer regions, other genes that are not oncogenes, or unidentified genomic elements (Unknown) or any combination thereof for LCINS and LCSS. **b)** Bar plot indicating the proportion of ecDNA containing a particular genomic element in LCINS vs LCSS. **c)** log_2_ median copy number (y-axis) for each class of ecDNA (x-axis) in LCINS and LCSS.

**Supplementary Fig 8. Highly sensitive pipeline for ecDNA detection that accounts for tumor purity and ploidy.**

**a)** Schematic depicting the methodology for the rescaled pipeline that considers tumor purity and ploidy. The logR track for the region containing the *EGFR* gene on chr7 and the purity and ploidy computed from Battenberg is indicated in one tumor as example (left). The exact location of the *EGFR* gene is shown by the dashed red lines. The equation that simultaneously accounts for sample purity and ploidy when computing the total copy number is shown, as well as the total copy number estimate when assuming a sample purity of 1.0 and ploidy of 2.0 (Uncorrected CN) *vs*. considering the purity and ploidy (Corrected CN). The Sashimi plot indicates the structural variants (colored arcs) and smoothed coverage around the *EGFR* region, which was annotated as ecDNA by AmpliconClassifier. **b)** Distribution of tumor purities for samples with an ecDNA annotation for both the default (left) and rescaled (right) pipelines. P-value for a Mann-Whitney test between ecDNA^+^ and ecDNA^-^ samples is indicated. The total number of samples in each group is also indicated. **c)** The ecDNA prevalence (y-axis), which is the proportion of ecDNA^+^ samples over the total number of samples, for all possible tumor purity cutoffs applied to samples (x-axis). The total number of samples available for analysis after each tumor purity cutoff is applied is also indicated. **d)** Gene expression (log_2_ CPM) of the EGFR gene for all samples with RNA-seq data available. LCINS samples with EGFR-ecDNA are indicated in red.

## SUPPLEMENTARY TABLE LEGENDS

**Supplementary Table 1. Co-occurrence of ecDNA genetic cargo in LCINS**

**Supplementary Table 2. Co-occurrence of ecDNA genetic cargo in LCSS**

**Supplementary Table 3. Recurrently amplified oncogenes on ecDNA in LCINS across unique genomic loci**

**Supplementary Table 4. Recurrently amplified oncogenes on ecDNA in LCSS across unique genomic loci**

**Supplementary Table 5. Additional *EGFR*-ecDNA detected by modified pipeline**

## Ethics declarations

Since the National Cancer Institute only received de-identified samples and data from collaborating centers, had no direct contact or interaction with the study participants, and did not use or generate identifiable private information, Sherlock-*Lung* has been determined to constitute “Not Human Subject Research (NHSR)” based on the federal Common Rule (45 CFR 46; https://www.ecfr.gov/cgi-bin/ECFR?page=browse).

## Data Availabiltiy

Normal and tumor-paired CRAM files for the WGS subjects of the Sherlock-Lung study and the EAGLE study have been deposited in dbGaP under the accession numbers phs001697.v2.p1 and phs002992.v1.p1, respectively.

## Code availability

All custom code developed for the analysis and generation of figures in this manuscript is publicly available in the GitHub repository (https://github.com/azhark2/Sherlock-Lung-ecDNA). Copy number variants were called using CNVkit (v.0.9.6) and rescaled for tumor purity and ploidy using estimates from Battenberg (v.2.2.9). Extrachromosomal DNA was detected and classified using AmpliconArchitect (v.1.3) and AmpliconClassifier (v.1.3.1). Chromothripsis was identified using ShatterSeek with structural variants called by Meerkat (v.0.189) and Manta (v.1.6.0). Telomere length was estimated using TelSeq (v.0.0.2). Mutational signatures were extracted using SigProfilerExtractor (v.1.1.3) with matrices generated by SigProfilerMatrixGenerator. Driver genes were identified using IntOGen (v.2020.02.0123). Additional publicly available software and algorithms used are detailed in the **Reporting Summary** and the **Methods** section. Any additional supporting scripts are available from the corresponding authors upon reasonable request.

## Author Contributions

Conceptualization, MTL, TZ, LBA; Methodology, AK, TZ; Formal Analysis, AK, TZ, MDG, JL, WZ, KB, EB, YY; Data Curation, TZ, PH, JM, CH, MM, OWL, MAN; Resources: NR, QL, DCW, LY, SJC, ACP, DC, KJ, AH, BH; Pathology evaluation: RH, MKB, LMS, PJ, CL, WDT, S-RY; Writing – Original Draft, AK; Writing – Editing, LBA, TZ, PHH, MTL; Writing – Review: All authors; Visualization: AK; Supervision, LBA, TZ, MTL.

## ACKNOWLEDGEMENTS

This research was supported by the Intramural Research Program of the National Institute of Health (NIH) (project ZIACP101231 to MTL), and by the Anne Wojcicki Foundation (Grant Number: LC009). The contributions of the NIH author(s) were made as part of their official duties as NIH federal employees, are in compliance with agency policy requirements, and are considered Works of the United States Government. However, the findings and conclusions presented in this paper are those of the author(s) and do not necessarily reflect the views of the NIH or the U.S. Department of Health and Human Services. We want to acknowledge the patients and the INCLIVA Biobank (PT17/0015/0049) integrated in the Spanish National Biobanks Network and in the Valencian Biobanking Network for their collaboration. This study was supported by the Health and Medical Research Fund of Hong Kong SAR, HMRF 03142856.

The related studies of Taiwan site were supported by grants from the Ministry of Health and Welfare, Taiwan DOH97-TD-G-111-026 (C.A.H.), DOH98-TD-G-111-015 (C.A.H.), DOH99-TD-G-111-028 (C.A.H.); DOH97-TD-G-111-029 (C.Y.C.), DOH98-TD-G-111-018 (C.Y.C.), DOH99-TD-G-111-015 (C.Y.C.), DOH97-TD-G-111-028(I.S.C.), DOH98-TD-G-111-017(I.S.C.), DOH99-TD-G-111-014(I.S.C.), and the Ministry of Science and Technology, Taiwan MOST109-2740-B-400-002 (C.A.H.), MOST110-2740-B-400-002 (C.A.H.), MOST1112740-B-400-002 (C.A.H.). This work has been supported in part by the Tissue Core at the H. Lee Moffitt Cancer Center & Research Institute, a comprehensive cancer center designated by the National Cancer Institute and funded in part by a Moffitt Cancer Center Support Grant (no. P30-CA076292). And, in part, by NIH (NCI) grant # U01CA209414 to the Boston Lung Cancer Survival Study of the Dana-Farber/ Harvard Cancer Center (D.C.C.). The authors would like to thank the team at the IUCPQ site of the Quebec Respiratory Health Network Biobank of the FRQS for their valuable assistance, and would like to thank the staff at Harvard University, Yale University, Roswell Park Cancer Institute and Roswell PI, Instituto Nacional de Cancerologia, Nice University Hospital Centre (Nice UHC) - University Côte d’Azur and the Nice Biobank CRB, Toronto University Health Network, and Mayo Clinic for their assistance providing samples and corresponding clinical data. The computational analyses reported in this manuscript have utilized the NIH high-performance Biowulf Cluster. We thank the study participants and the staff at Westat Inc. for their assistance in collecting samples and corresponding clinical data. Part of this work was supported by the US National Institute of Health grants R01ES032547, R01CA269919, P01CA281819, and U01CA290479 to LBA as well as by LBA’s Packard Fellowship for Science and Engineering. MD-G was awarded with a fellowship within the “Generación D” initiative, Red.es, Ministerio para la Transformación Digital y de la Función Pública, for talent attraction (C005/24-ED CV1), funded by the European Union NextGeneration EU funds, through PRTR. The research in this study was also supported by UC San Diego Sanford Stem Cell Institute. This work utilized the computational resources of the NIH HPC Biowulf cluster (https://hpc.nih.gov). The funders had no roles in study design, data collection and analysis, decision to publish, or preparation of the manuscript.

