## Supplementary Figures for "Examining the Role of Extrachromosomal DNA in Lung Cancer"

**Supplementary Fig. 1**

**a**

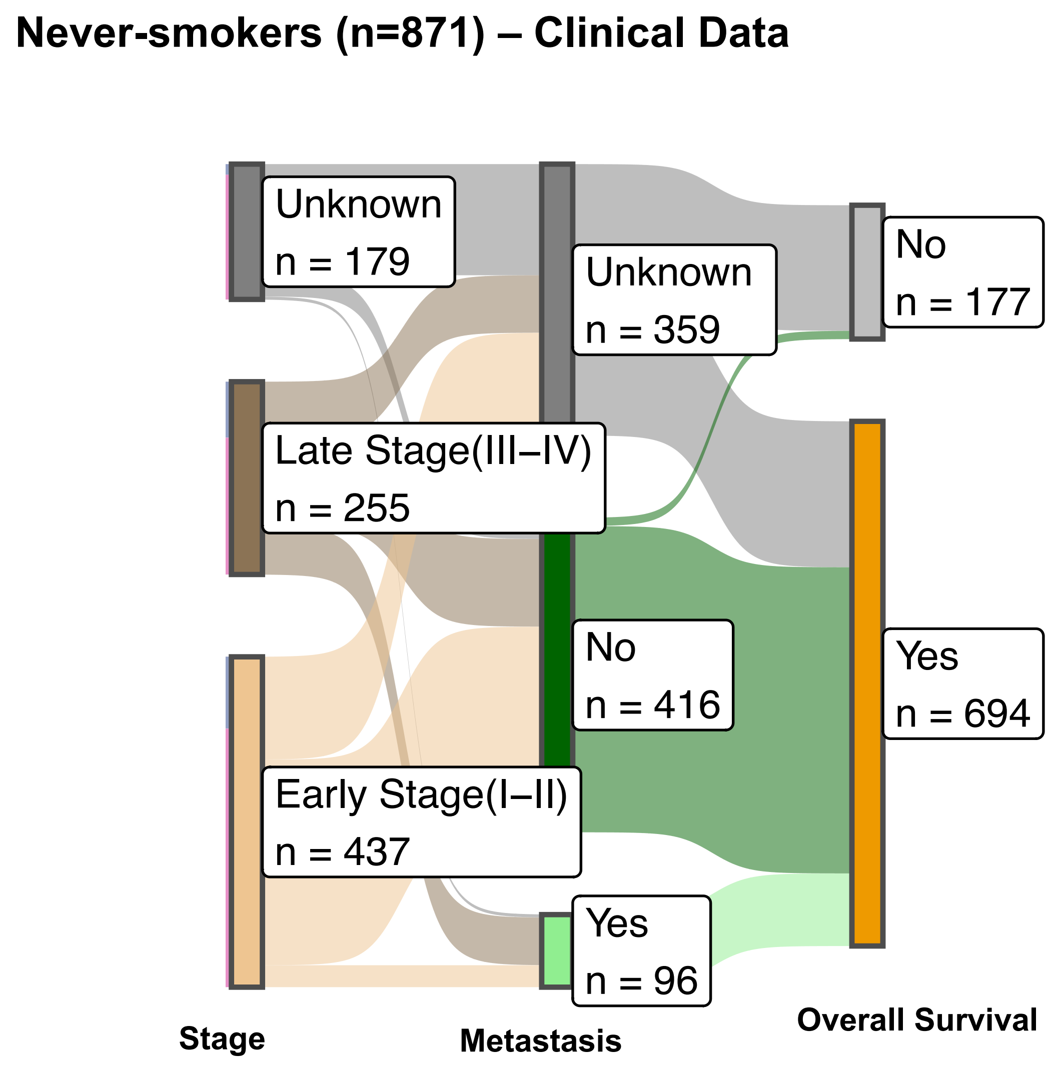


**b**

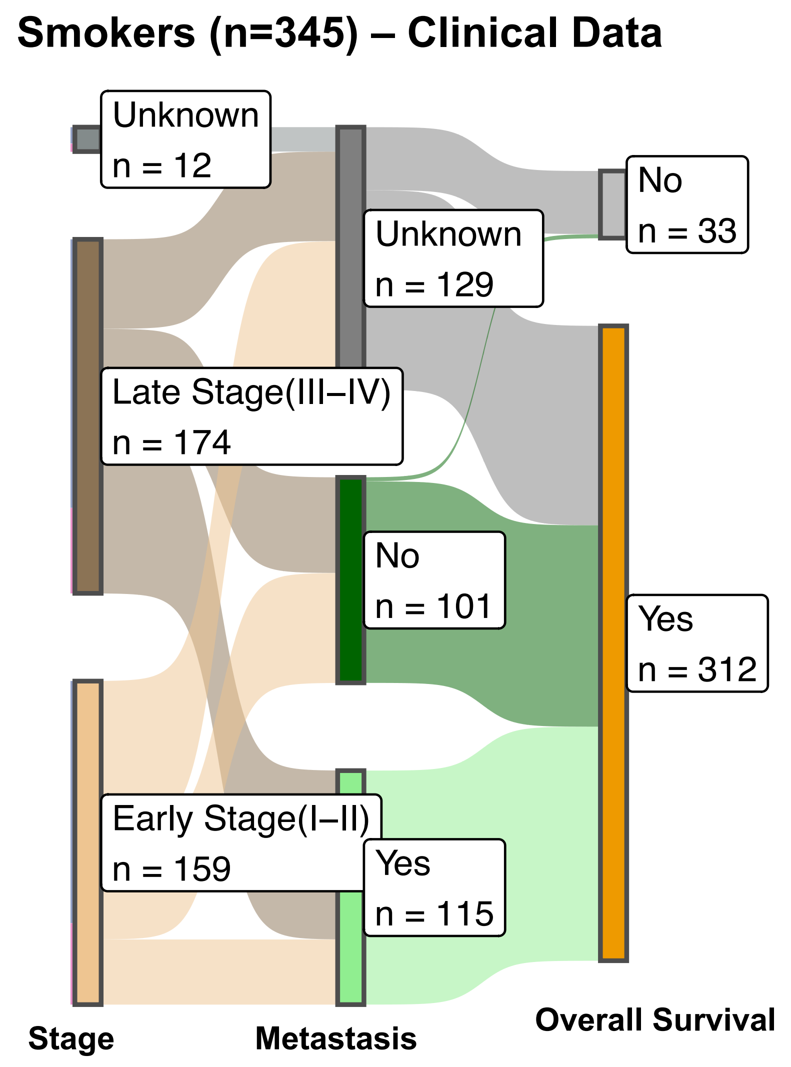

Supplementary Fig. 2

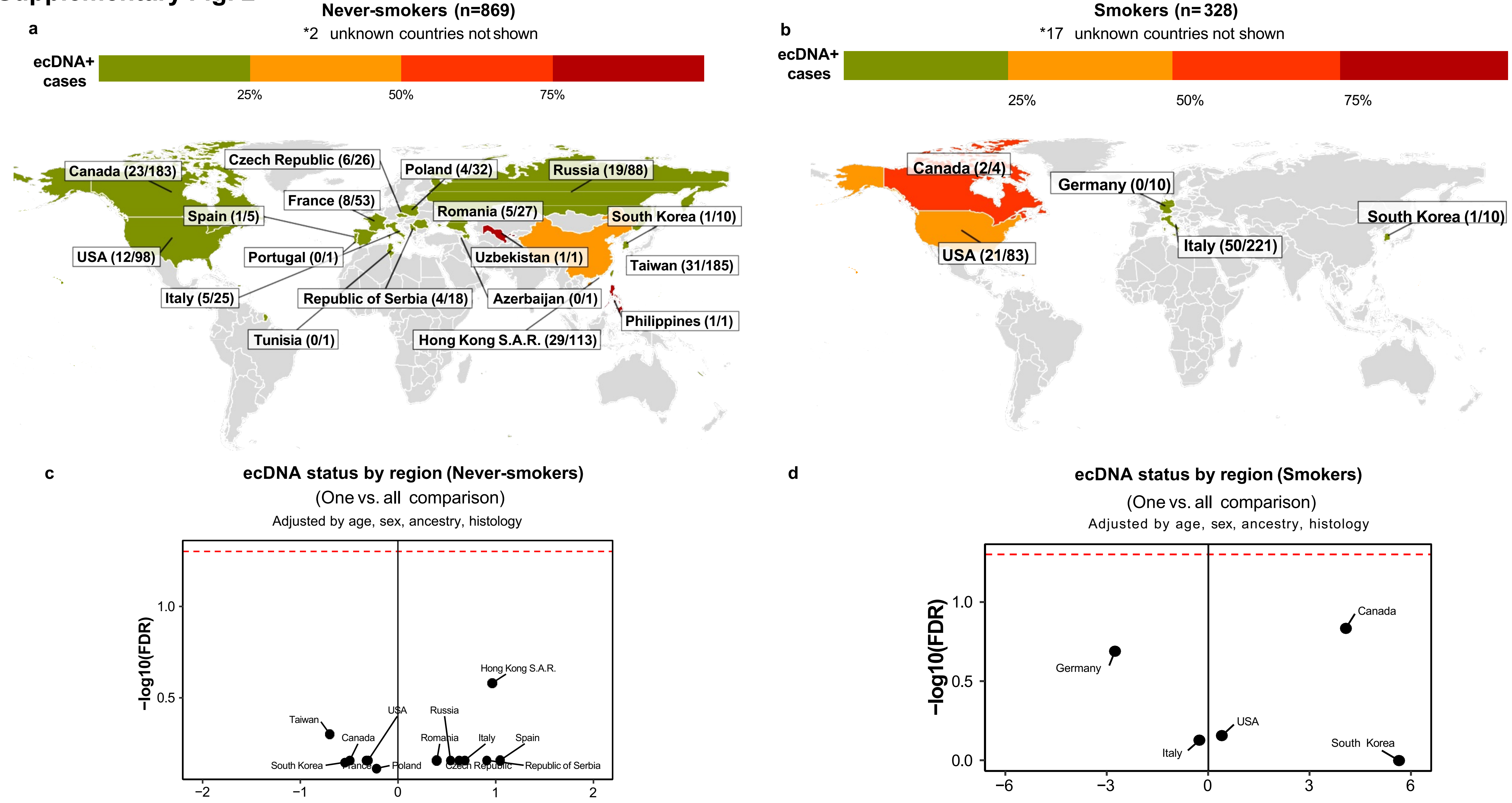

**Supplementary Fig. 3**

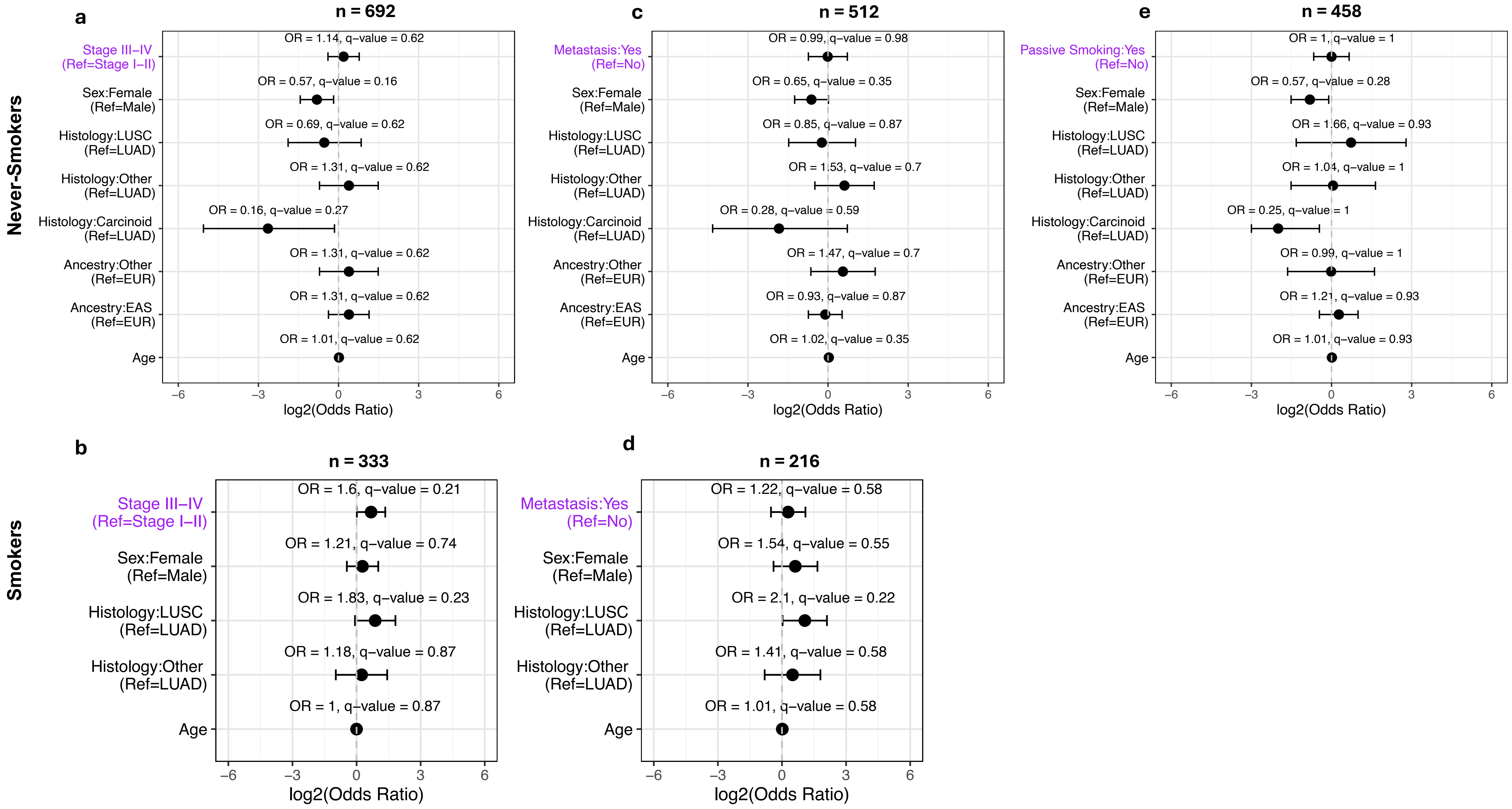

**Supplementary Fig. 4**

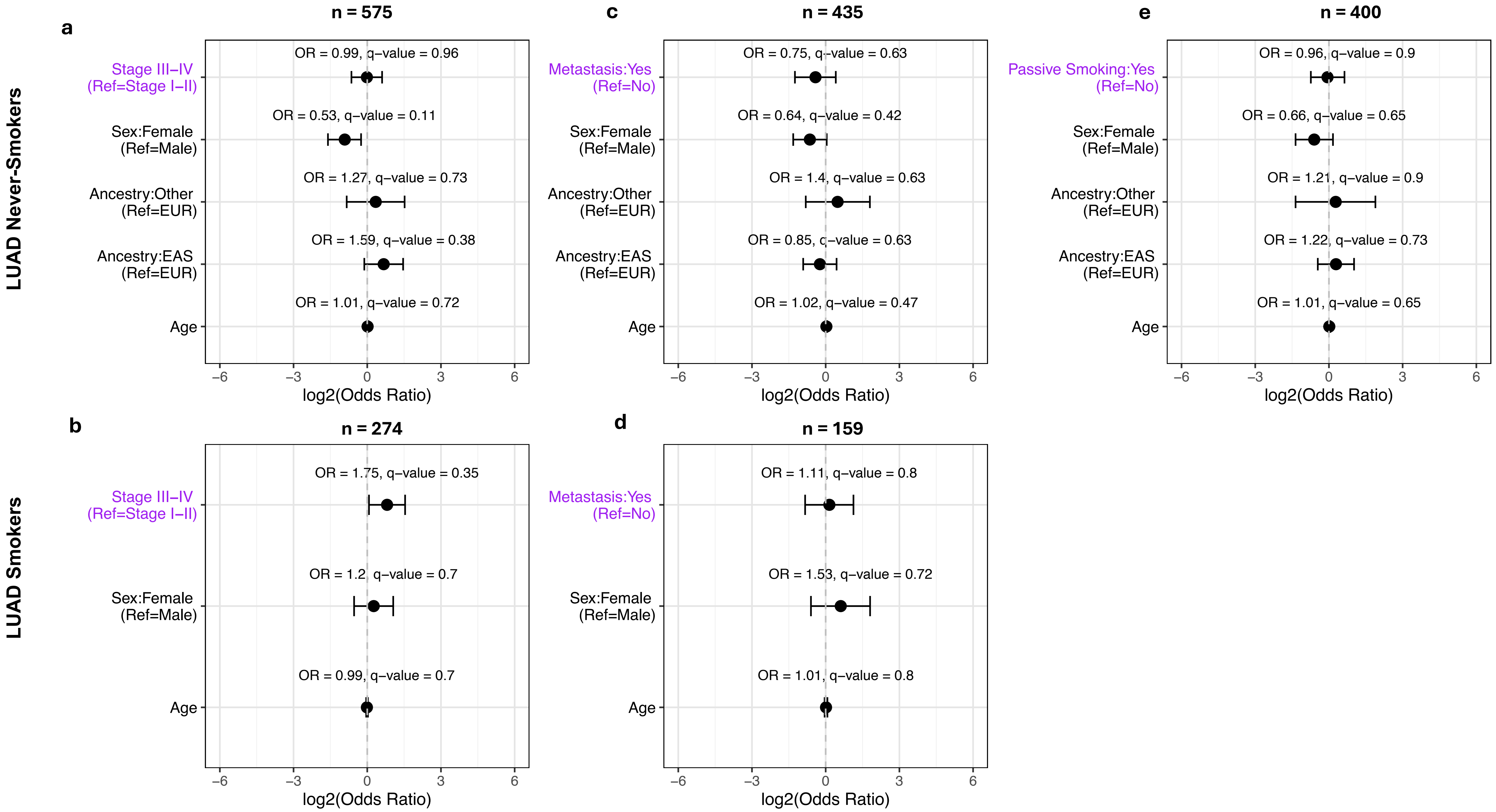

### Supplementary Fig. 5

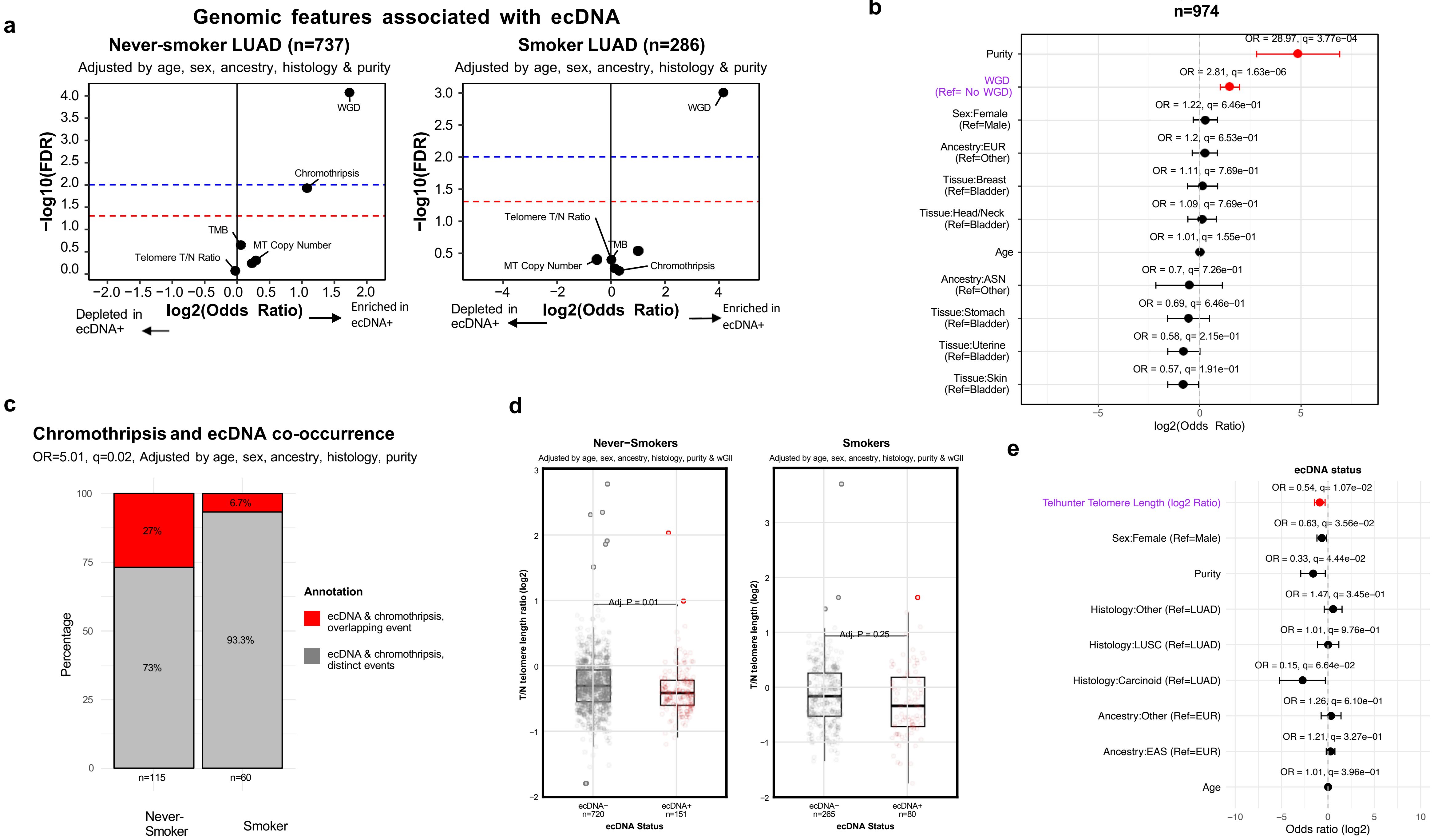

#### Supplementary Fig. 6

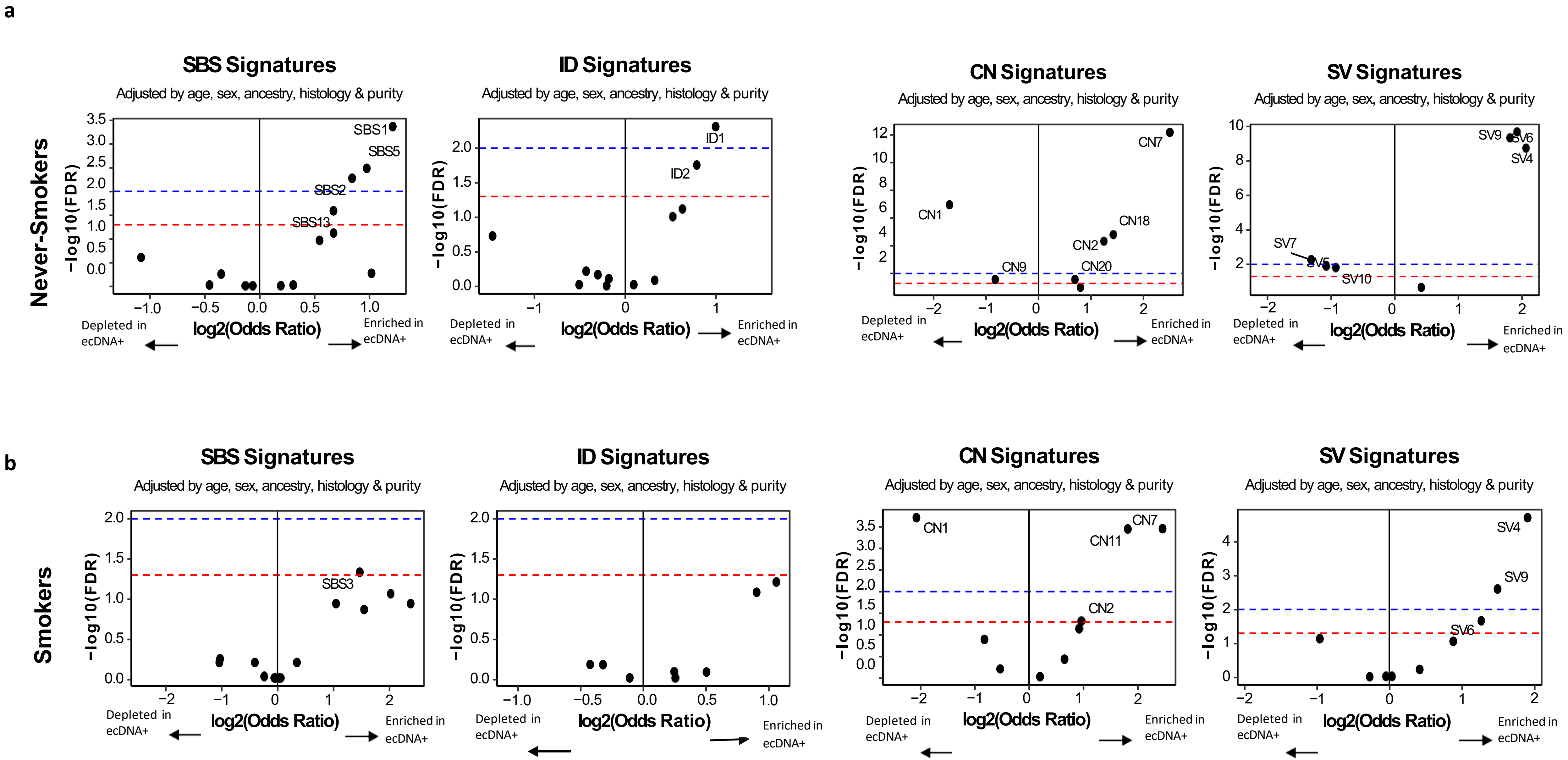

### Supplementary Fig. 7

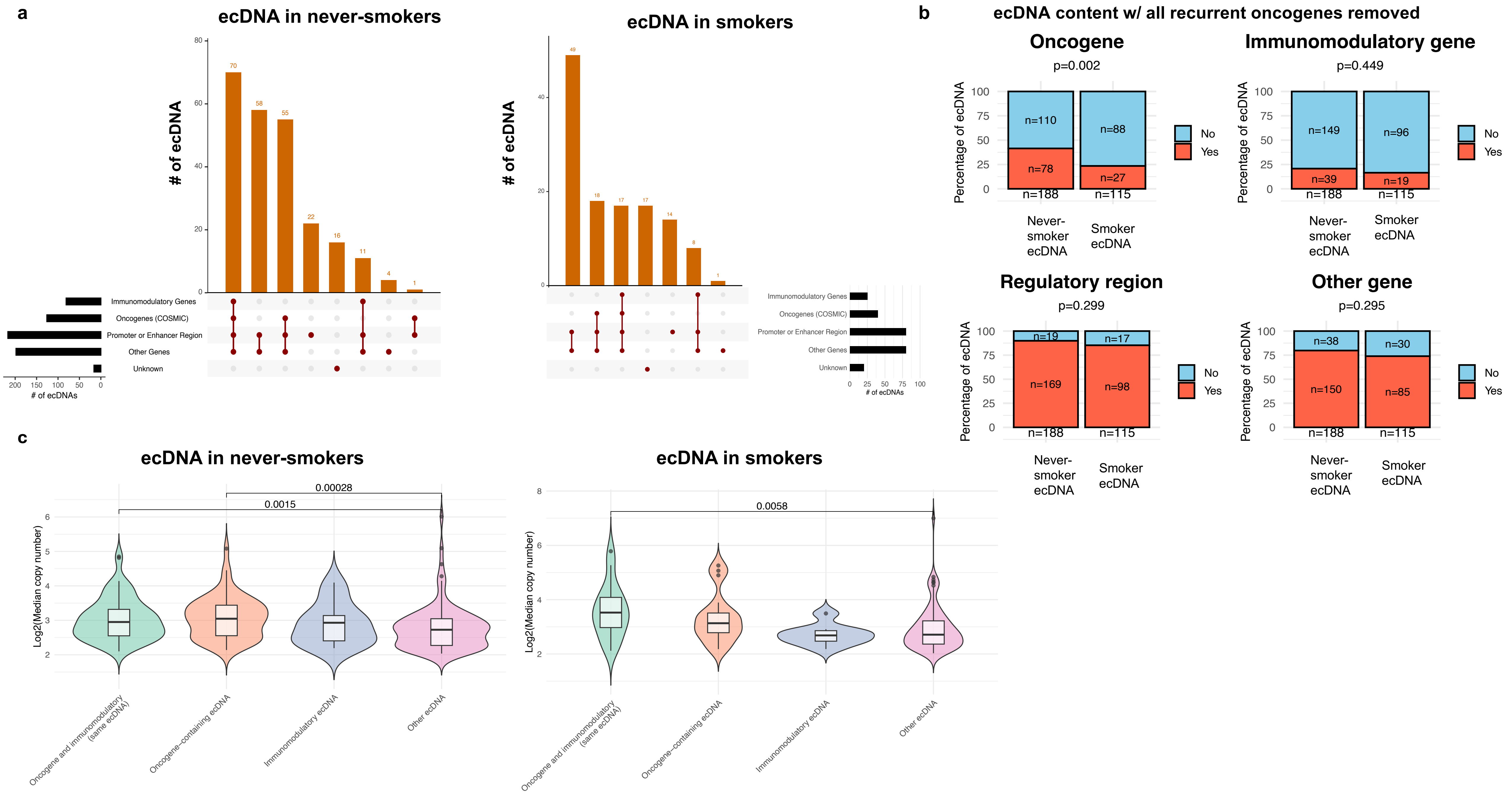

#### Supplementary Fig. 8

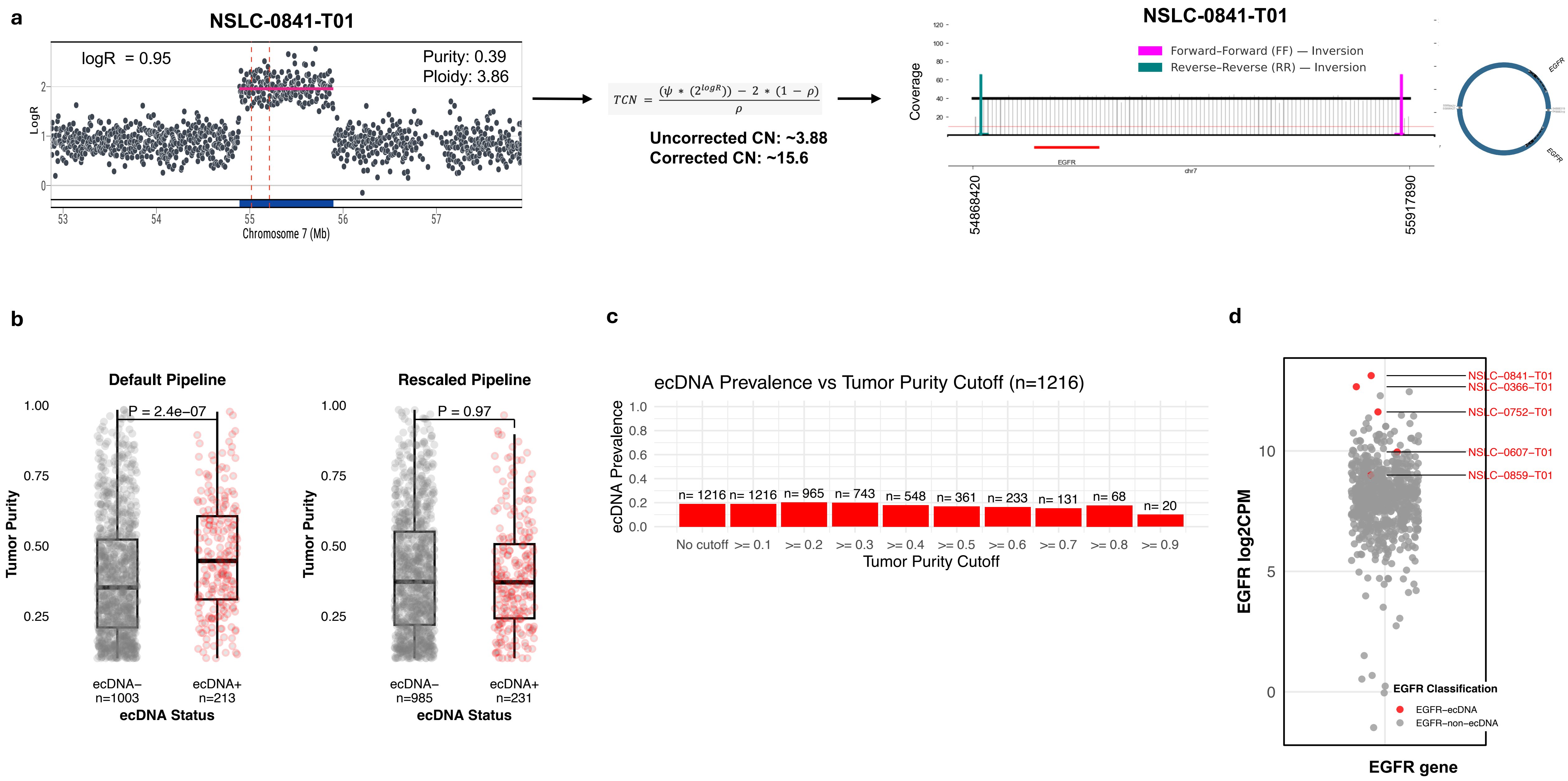
